# Genomic blueprints of a dynamic deep sea: contrasting adaptive strategies potentially drive seasonal microbial succession

**DOI:** 10.64898/2026.11.16.767118

**Authors:** Binghe Zhao, Alexandru S. Barcon, Song Zhong, Yinghui He, Yong Wang

## Abstract

Genetic variation within marine microbial populations can help reveal how deepsea communities respond to environmental change. However, population-level genomic variation in deep pelagic waters remains poorly understood. Here, we analyzed metagenomic and metatranscriptomic datasets from ∼1,080 m depth in the South China Sea to compare genomic variation between summer and winter microbial assemblages. Among 11 persistent metagenome-assembled genomes (MAGs) detected in both summer and winter samples, we observed differences in abundance, nucleotide diversity (π), pN/pS ratios, and gene-level differentiation. Two enriched populations showed contrasting patterns, with *Sphingobium yanoikuyae* enriched in winter samples and *Methyloversatilis universalis* enriched in summer samples. The winter-enriched *S. yanoikuyae* population carried winter-associated accessory genes related to restriction-modification and toxin-antitoxin systems, consistent with a defense-oriented persistence profile. In contrast, the summer-enriched *M. universalis* population showed higher nucleotide diversity in summer samples and divergence in genes related to motility, transport, and cobalamin acquisition. Across persistent MAGs, low pN/pS ratios indicated predominantly purifying selection, whereas gene-specific divergence and flexible accessory gene content suggested population-level adaptive potential under contrasting winter and summer conditions. These results reveal population-genomic differences associated with winter-summer microbial shifts in the deep sea.

**Importance:** The deep ocean is a vast ecosystem where microorganisms experience environmental changes that can influence their survival and ecological roles. However, how deep-sea microbial populations respond to contrasting environmental conditions at the genetic level remains poorly understood. By combining genomic and gene expression analyses, this study reveals distinct population-level responses of two deepsea bacteria, *Methyloversatilis universalis* and *Sphingobium yanoikuyae*, under winter and summer conditions in the South China Sea. The summer-enriched *M. universalis* showed greater genetic diversity and differences in genes associated with motility, nutrient acquisition, and environmental sensing, whereas winter-enriched *S. yanoikuyae* carried genes associated with cellular defense and persistence. These contrasting patterns suggest that deep-sea microorganisms may employ different strategies to maintain their populations under changing environmental conditions. Our findings extend the understanding of microbial adaptation in the deep ocean beyond changes in community composition to include variation within microbial populations.

## Introduction

The deep sea, comprising the vast water column below the euphotic zone, is the largest yet least explored biome on earth^1^. It plays a major role in global biogeochemical cycles by acting as a long-term reservoir for carbon and nutrients sequestered from the surface^1,2^. Exploring the dynamics of marine microorganisms under fluctuating physicochemical conditions is crucial for understanding future marine ecosystems^3^. For example, long-term studies of ocean surface waters have revealed the repetitive seasonal and annual variations in temporal dynamics^4^. Recent time-series studies leveraging 16S rRNA gene amplicon sequencing have revealed that mesopelagic microbial communities exhibit pronounced seasonal shifts in composition and diversity, probably coupled to surface phytoplankton dynamics and particle flux. In particular, microbial assemblages at 500-m depth have been shown to mirror surface seasonal patterns, with community turnover driven by both the downward transport of bloom-associated taxa and winter convective mixing that facilitates vertical exchange of endemic deep-sea lineages^5,6^. However, the adaptive mechanisms of marine microorganisms under different seasonal conditions remain unclear.

To understand dynamics of microbial populations in the fluctuating deep-sea environment, we must resolve the fitness processes acting at the level of their genomes. The population genomics has provided profound insights into surface-ocean adaptation by revealing fine-scale genetic variations (i. e. single nucleotide polymorphism; SNP) of discrete ecotypes in *Prochlorococcus* populations^7^, horizontal gene transfer and selective sweeps among *Vibrio* populations colonizing chitin particles^8^, and genome streamlining that shapes the ecology of the ubiquitous *Pelagibacter*^9^. High rates of nonsynonymous to synonymous substitutions (pN/pS) of positively selected genes have been reported to predict protein translocation, DNA synthesis and motility in some deep-sea bacteria^10^.Despite its ability to resolve selection pressures, nucleotide diversity, recombination rates and pangenome dynamics, this approach has rarely been applied to deep pelagic microbial populations sampled under contrasting winter and summer conditions. Furthermore, metagenomic studies have begun to catalog genomic variations in deep-sea sediments^11^, hydrothermal vents^12^, and subseafloor aquifers^13^; however, the temporally resolved, seasonal sampling required to capture adaptive evolution in action remains to be resolved.

The South China Sea (SCS) is one of the largest marginal seas^14^. The strong northeast monsoon of winter SCS drives intense vertical mixing, deepens the mixed layer, and facilitates the downward transport of nutrients, lithogenic material, and trace metals from polluted rivers, as evidenced by seasonal shifts in water column structure and biogeochemical fluxes^15–17^. Conversely, the summer is characterized by strong thermal stratification, shallower mixed layers, and variable surface productivity^18,19^. This pronounced environmental oscillation creates a potent selective landscape, making SCS an ideal setting to investigate the genomic basis of microbial adaptation to seasonal change. Our previous *in situ* metatranscriptomic and metagenomic study at ∼1,080 m depth revealed differences in active microbial community structure and metabolism between winter and summer samples^20,21^.

In this study, we utilized the high-resolution, winter and summer meta-omics datasets from four ∼1,080-m stations in SCS to investigate population-level genomic variation. By reconstructing metagenome-assembled genomes (MAGs) and tracking their population-level dynamics across contrasting winter and summer regimes, we focused on 11 persistent species with seasonal succession and transcriptional difference, providing insights into evolutionary processes in the dynamic deep sea. Notably, summer-enriched *M. universalis* population, exhibited higher nucleotide diversity and gene-specific divergence during their peak abundance, particularly in genes associated with cell motility. We therefore hypothesized that seasonal dominance may be associated with expansion of genetically heterogeneous populations and gene-level diversification in niche-related functions.

## Results

### Deep-sea microbial communities exhibit pronounced seasonal shifts in composition

To improve understanding in seasonal intraspecific variation of deep-sea microorganisms at population level, we performed the analysis on a set of MAGs that were reconstructed from the metagenomes for 18 deep-sea time-series water samples covering winter (n = 12) and summer (n = 6) of SCS obtained by the Multiple in situ Nucleic Acid Collections apparatus (MISNAC)^22^(Figs 1a and S1). This set includes 1,175 MAGs that were clustered at 95% average nucleotide identity (ANI) into 423 prokaryotic MAGs at species level^20^. To ensure the robustness of the mapping results and minimize potential interference from reads originating from non-target organisms, stringent criteria were applied for read recruitment. Reads sharing >95% identity with the respective MAGs were retained. Among these MAGs with an average coverage >10 X and a breadth of coverage >0.5 (i.e., >50% of the genome length was covered by reads) within a sample were selected for downstream analysis. This resulted in a subset of 11 MAGs with sufficiently high level and breadth of coverage by the metagenomic reads between the two seasons (Supplementary Table 1).

**Fig 1.**
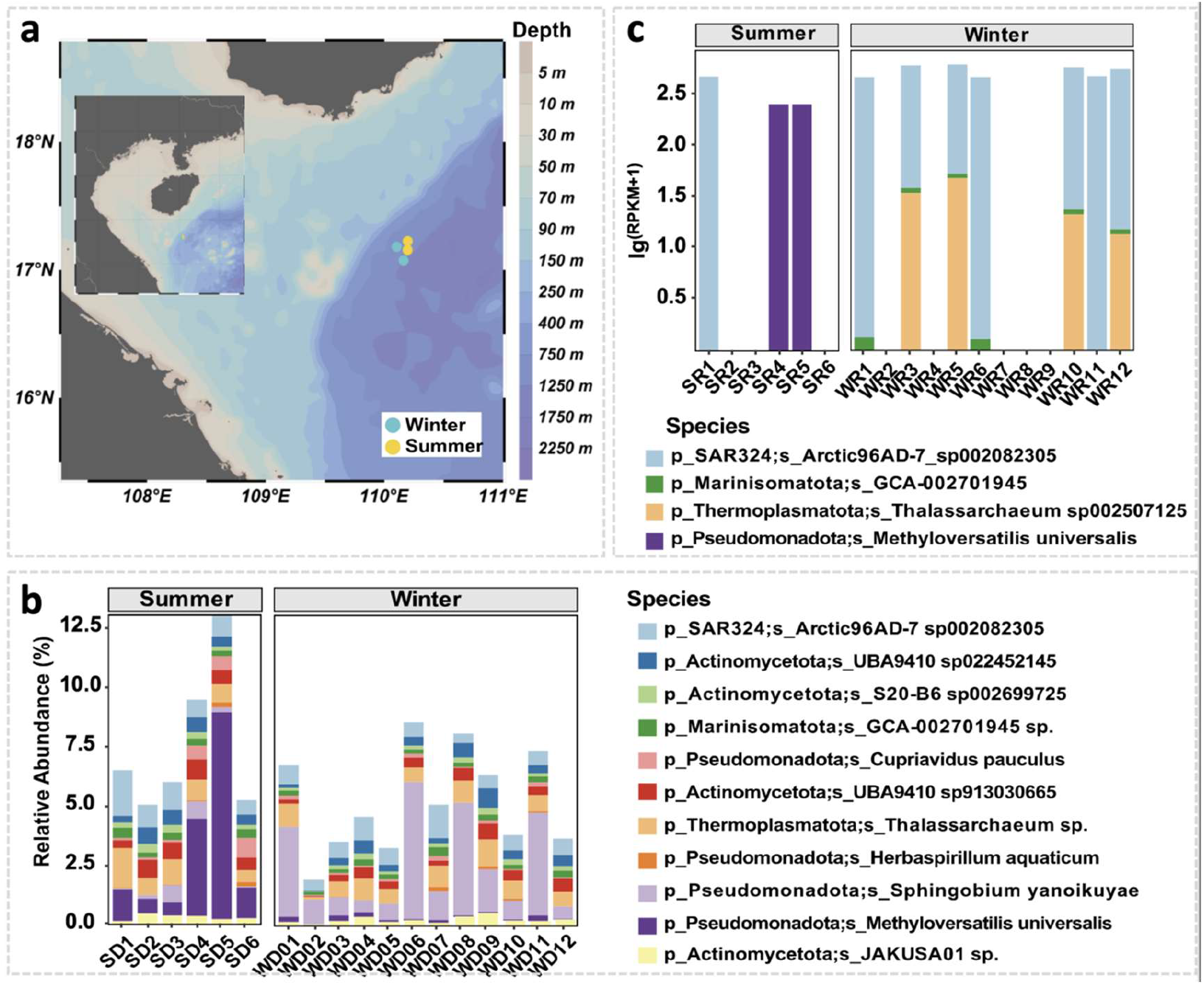
Seasonal dynamics of the deep-sea microbial communities in the South China Sea. (a) Map of the sampling stations at 1,000-1,100-m depth. Color shading indicates bathymetry. (b) Relative abundance (%) of the 11 most persistent bacterial genera across 18 metagenomics datasets. Summer samples (SD; n = 6) and winter samples (WD; n = 12) show clear seasonal succession, with *M. universalis* (violet) dominant in summer and *S. yanoikuyae* (light purple) enriched in winter. (c) Absolute abundance (RPKM) based on coverage of metatranscriptomic reads on the MAGs of the same species across seasons.

Seasonal environmental variations exerted a pronounced influence on microbial community composition. The deep-sea water was dominated by the species affiliated with Alphaproteobacteria, Gammaproteobacteria and SAR406, more or less displaying transcriptional activity (Fig. S2). The 11 MAGs included in the species-level analysis exhibited distinct seasonal shifts in relative abundance (Fig. 1b). For instance, the Wilcoxon rank-sum test confirmed significant seasonal differences for 6 of the 11 species (*p* < 0.05) (Supplementary Table 2); The *M. universalis* and *S. yanoikuyae* showed the pronounced seasonal contrasts (*p* < 0.005). By carrying on the fold change analysis of Reads Per Kilobase per Million mapped reads (RPKM) – based abundance (average RPKM value of winter samples divided by average RPKM of the summer samples) revealed that *M. universalis* exhibited a strong preference for summer (ratio = 0.05), whereas *S. yanoikuyae* showed a strong preference for winter (ratio = 5.75; Supplementary Table 2). Metatranscriptomic analysis provided further insights into the seasonal activity patterns of the microbial community. We observed that *M. universalis* was transcriptionally active exclusively in summer, which aligns well with its strong seasonal preference and dominance during this period. In contrast, transcripts for *S. yanoikuyae* were not detected in either season (Fig. 1c), suggesting a lack of metabolic activity at the sampling depth despite its established genomic presence and winter enrichment. Beyond these dominant species, Arectic96AD-7 of SAR324 demonstrated transcriptional activity across both seasons. Additionally, *Thalassarchaeum* showed greater transcriptional activity in winter, despite exhibiting significantly higher genomic abundance in summer (*p* < 0.05) (Fig. 1c).

### Seasonal clonal expansion and SNP dynamics in dominant microbial populations

To assess whether seasonal abundance shifts were associated with clonal expansion, we examined temporal abundance trajectories of the 11 MAGs across all 18 samples (Fig. 2a). Both *S. yanoikuyae* and *M. universalis* displayed non-uniform, pulsed growth patterns, characterized by abrupt, high-magnitude increases at specific sampling time intervals. For example, *M. universalis* exhibited the highest RPKM 61.57 in sample SD5, which was 5.3 times greater than the subsequent sample SD6 and 2.1 times greater than the previous sample SD4. Similarly, *S. yanoikuyae* reached its peak RPKM value of 36.67 in the WD6 sample. This level was 3.7 times higher than at the next time point WD7 and 5.9 times higher than at the preceding time point WD5, remaining relatively low during other periods (Fig. 2a). SNPs were detected in the 11 shared MAGs by mapping the time-series metagenomics reads of the winter and summer samples on the MAGs (Fig. 2a).

**Fig. 2.**
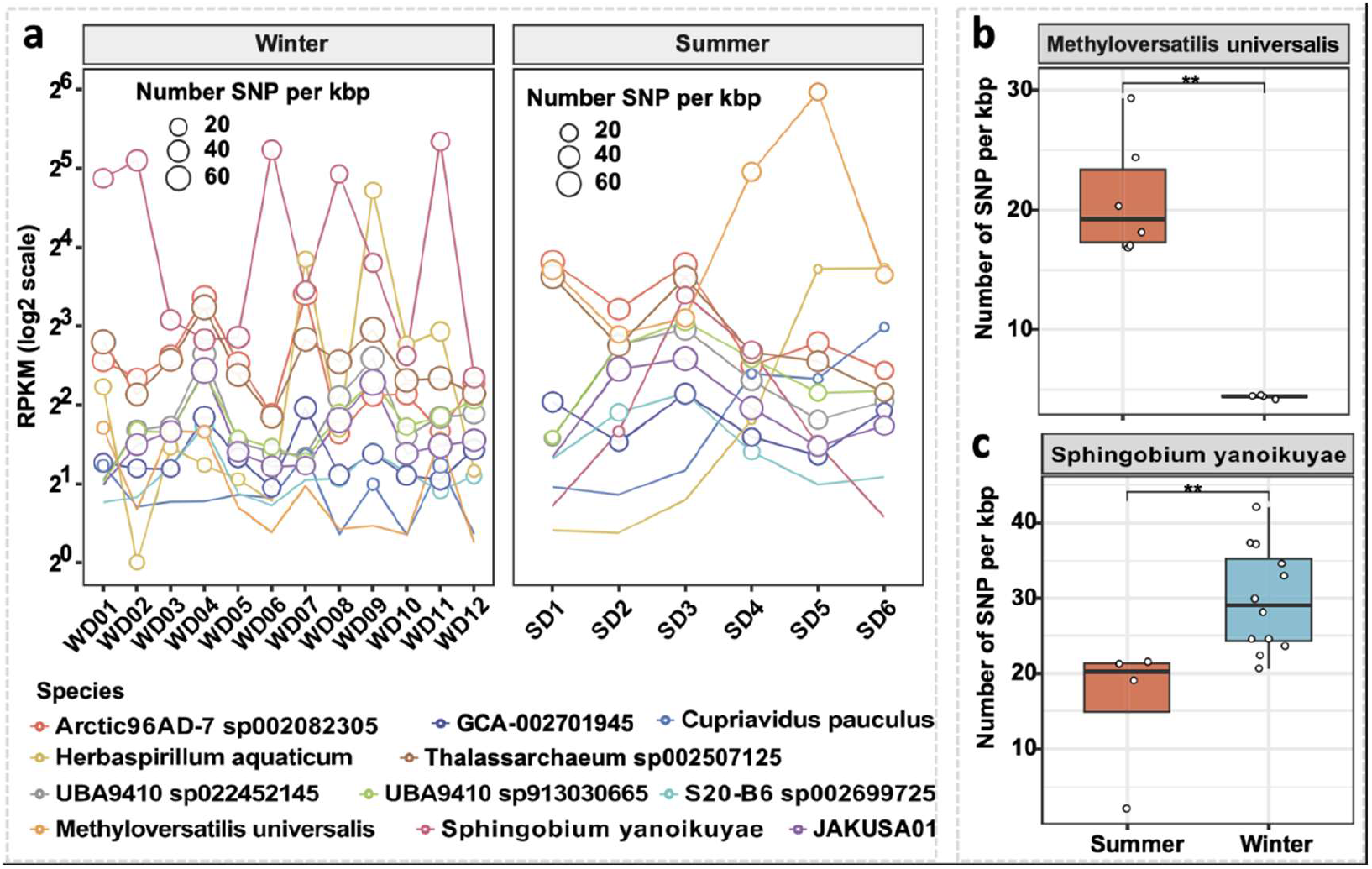
Seasonal abundance dynamics and SNP density of dominant populations. (a) Abundance of the 11 shared MAGs across all samples (RPKM). Circle size indicates SNP density (SNPs per kbp). (b, c) Boxplot comparison of SNP densities between summer and winter for the two species. Asterisks (**) indicate significant differences (Wilcoxon rank-sum test, p < 0.01).

The relationship between abundance and genomic diversity showed that high-abundance samples frequently coincided with high SNP density (Fig. S6). The decline of *S. yanoikuyae* abundance from winter to summer contrasted sharply with the opposing trend of *M. universalis*. We quantified the number of detected SNPs, and showed that both genera accumulated significantly more SNPs in their respective season of dominance (*p* < 0.001, Wilcoxon rank-sum test). *S. yanoikuyae* carried a mean of 159,966 ± 37,866 SNPs in winter versus 85,844 ± 49,889 SNPs in summer. In contrast, *M. universalis* carried 79,043 ± 18,611 SNPs in summer (n = 6) but only 16,588 ± 584 SNPs in winter (n = 4). We further compared SNP density between seasons for the two dominant populations. The results showed that *M. universalis* had significantly higher SNP density in summer than in winter, whereas *S. yanoikuyae* displayed the reverse pattern (*p* < 0.01; Fig. 2b, c). Additionally, the *Herbaspirillum* and *Thalassarchaeum* MAGs were characterized with significant high SNP density in the winter samples, although significant differences in abundance have not been observed across summer and winter (Figs S3, S4).

### Persistent populations display season-dependent genomic diversity and structure

We quantified nucleotide diversity (π) and nonsynonymous-to-synonymous polymorphism ratios (pN/pS) across the 11 MAGs detected in both seasons. As an indicator of conservative degree of a gene, high overall pN/pS ratios of the genes in a species imply rapid evolution to adapt to a new environment^23^. The Arctic96AD-7 MAG showed the highest average pN/pS value, while the *S. yanoikuyae* MAG displayed the lowest on average. The *S. yanoikuyae* and *M. universalis* MAGs exhibited low π (0.0108 ± 0.0039 and 0.0076 ± 0.0034, respectively) (Fig. 3a). To assess how seasonal environmental variations affected these selection pressure and diversity metrics, we compared the seasonal difference of π and pN/pS across these metrics. The result showed that the *M. universalis* had significantly higher π in summer (*p* < 0.01), corresponding to the season when its abundance and transcriptional activity were the highest (Fig. 1c, d). However, the *S. yanoikuyae* displayed relatively stable diversity between seasons (*p* > 0.05), regardless of seasonal difference in population size. *Thalassarchaeum* sp. transcriptionally active in winter showed significant lower pN/pS value but higher nucleotide diversity in winter, which indicates a strong purifying selection force against nonsynonymous mutations (Fig 1c, Supplementary Table 3). Across the 11 shared MAGs, four showed significant seasonal differences in π, and six showed significant differences in pN/pS (Supplementary Table 3). Additionally, the π versus pN/pS scatter plot (Fig. 3a) depicted the diverse patterns in genomic diversity and selection across all 11 MAGs, with detailed seasonal differences presented in Supplementary Table 3.

**Fig. 3.**
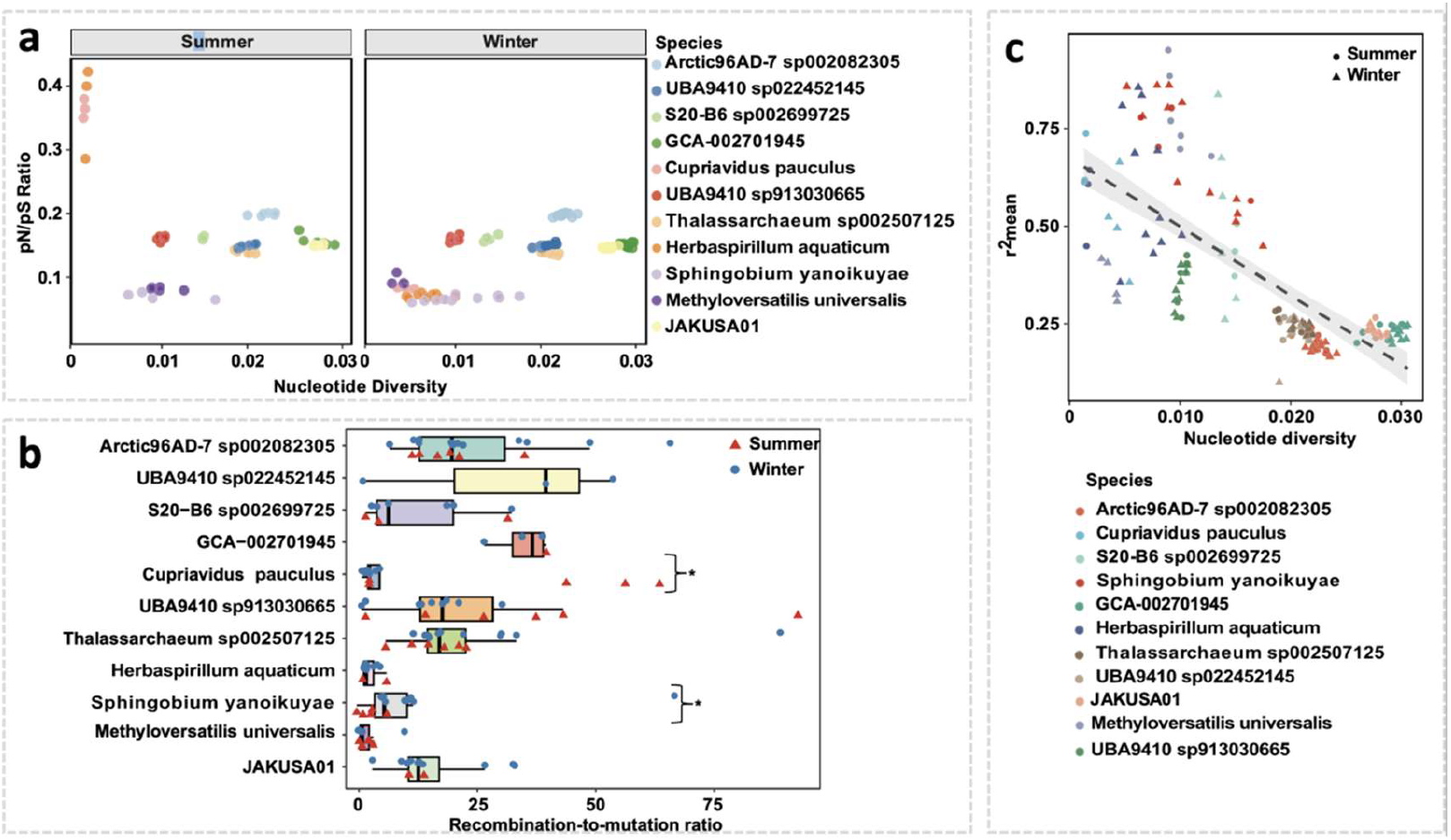
Genomic diversity and selection pressures of persistent deep-sea prokaryotic populations. (a) Scatter plot of nonsynonymous-to-synonymous polymorphism ratios (pN/pS) versus nucleotide diversity (π) across 11 shared MAGs. Circles represent summer samples and triangles represent winter samples. Colors indicate individual MAGs, showing distinct clustering by taxon and season. Overall low pN/pS values indicate purifying selection, while nucleotide diversity varies among taxa. (b) Recombination- to-mutation ratio (gamma/mu) calculated for the 11 populations. Statistically significant differences between the gamma/mu values of the two seasons are noted with asterisks (p < 0.05). (c) Scatter plot of nucleotide diversity (π) versus mean linkage disequilibrium (r²) for the same MAGs, with circles representing summer samples and triangles representing winter samples. Colors indicate individual MAGs. A strong negative correlation was observed (Spearman’s ρ = -0.808, p < 0.001), with the species M. universalis showing higher nucleotide diversity in summer compared with winter.

Furthermore, *Cupriavidus pauculus* showed a significant increase in pN/pS from a low winter value (0.097) to a relatively high summer value (0.269) (*p* < 0.001), while its π values remained low and stable across seasons (winter: 0.005; summer: 0.002). *Herbaspirillum aquaticum* exhibited a significant rise in pN/pS from a low winter value (0.075) to a relatively high summer value (0.264) (*p* < 0.001) and a significant decrease in π from a moderate winter value (0.007) to a low summer value (0.002) (*p* < 0.01). GCA-002701945 displayed a significant increase in pN/pS from an intermediate winter value (0.154) to a slightly higher summer value (0.161) (*p* < 0.05), coupled with a significant decrease in π from a high winter value (0.030) to a slightly lower summer value (0.028) (p < 0.05). *JAKUSA01* showed a significant increase in pN/pS from an intermediate winter value (0.150) to a slightly higher summer value (0.154) (p < 0.001), but its π values remained relatively high and stable across seasons (winter: 0.027; summer: 0.028). Conversely, S20-B6 sp002699725, UBA9410 sp022452145, and UBA9410 sp913030665 maintained stable pN/pS values and π values across seasons, with no significant seasonal differences observed for either metric (Fig. 3a, Supplementary Table 3).

Building on these patterns in diversity and selection, we assessed the role of recombination in shaping population structure. Both *S. yanoikuyae* and *M. universalis* exhibited a decay in linkage disequilibrium with increasing SNP distance (Fig. S5), likely due to frequent recombination events. The analysis of inStrain-derived r² values^24^ across all 11 MAGs showed significant variation between taxa and a strong negative correlation with nucleotide diversity (Spearman’s *ρ* = -0.808, *p* < 0.001) (Fig. 3c). In addition, we further estimated the ratio of the recombination rate to mutation rate (gamma/mu) for these 11 populations, finding that this value varied significantly across populations (Supplementary Table 4). GCA-002701945 exhibited the highest ratio at 35.91, whereas *S. yanoikuyae* and *M. universalis* showed relatively lower rates of 10.95 and 2.57, respectively. For *Cupriavidus* and *S. yanoikuyae* populations, we detected statistically significant differences in gamma/mu values (*p* < 0.05) between the summer and winter samples. Specifically, *Cupriavidus* populations showed higher recombination rates in summer (34.67) compared to winter (3.01), and *S. yanoikuyae* populations exhibited higher recombination rates in winter (16.48) than in summer (3.16) (Fig 3b). This suggests the possibility of season-specific environmental influences on the recombination rate of populations within the same species. Given that r² quantifies linkage disequilibrium (LD) between single-nucleotide variants^25^, lower r² values at higher nucleotide diversity are consistent with more frequent recombination events. However, the *M. universalis* MAG deviates from this trend by exhibiting both higher nucleotide diversity and higher r² values in summer compared to winter.

### Gene-level analysis reveals signatures of seasonal adaptation

We examined gene-level differentiation between seasons for the two dominant species, *S. yanoikuyae* and *M. universalis*, by computing gene-wise fixation index F_ST_ for all genes shared between the winter and summer populations^26^, followed by z-score normalization to identify outliers. Across all shared genes, mean F_ST_ was 0.07 in the species *S. yanoikuyae* and 0.31 in *M. universalis*. We detected 39 high-F_ST_ genes in the genus *M. universalis* with values up to 0.83, and high-F_ST_ genes up to 0.54 in the genus *S. yanoikuyae* (*p* < 0.05, right-tailed test; Fig. 4a, b).

**Fig 4.**
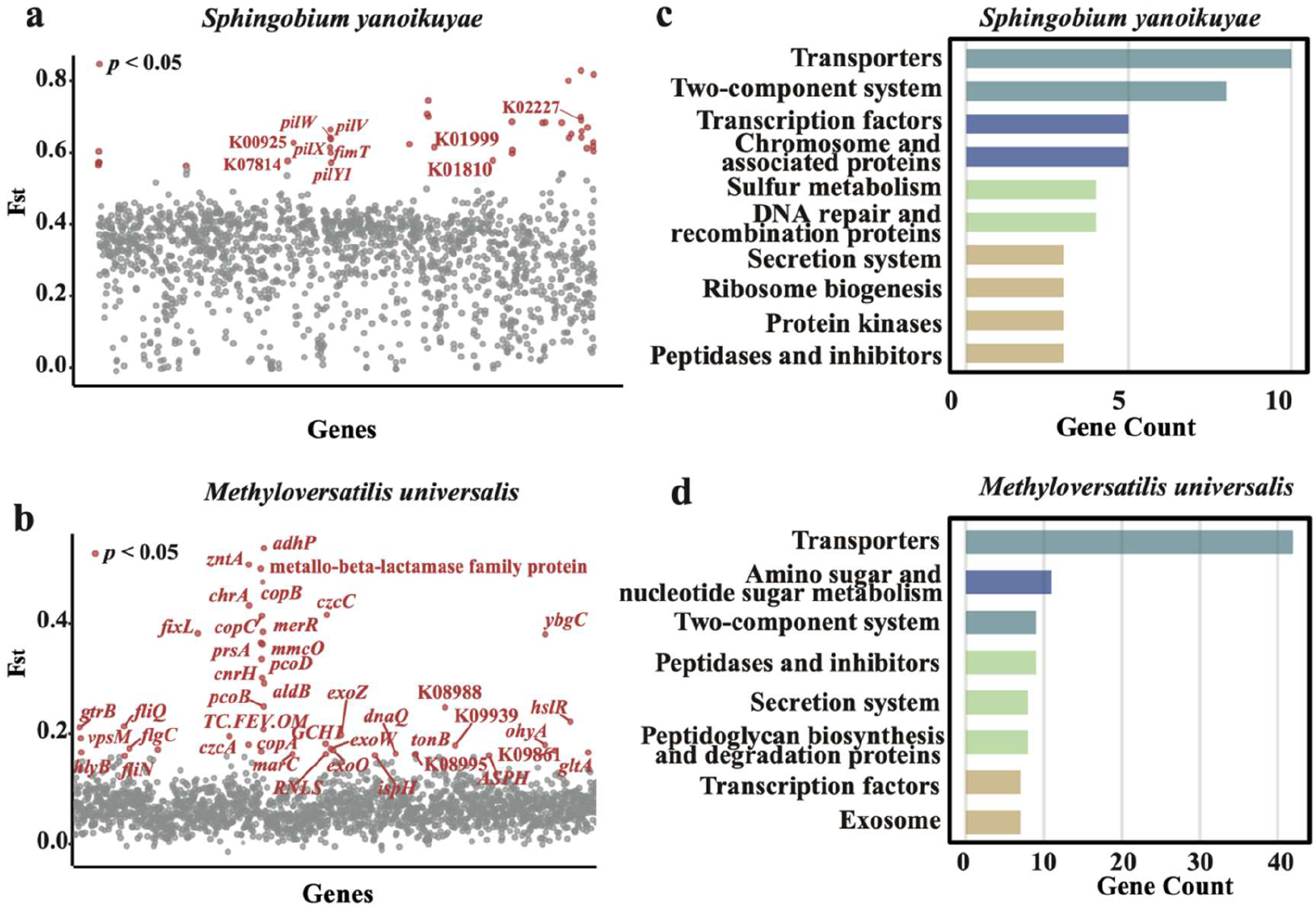
KEGG pathway enrichment of genes with significant seasonal differences in SNP density. Differentiated genes (marked in red) with elevated F_ST_ values with statistical significance (adjusted p value of < 0.05 [by a Z test]) from a (S. yanoikuyae) and b (M. universalis). Abbreviations for putative gene functions are as follows: pilW, pilV, pilX, pilY1 type IV pilus assembly protein; fimT, type IV fimbrial biogenesis protein; fliQ, flagellar biosynthesis protein; vpsM, polysaccharide biosynthesis protein; fliN, flagellar motor switch protein; flgC, flagellar basal-body rod protein; MIP, MIP channel protein; exoZ, exopolysaccharide production protein; exoW, exoO, succinoglycan biosynthesis protein.(c) Genes with higher SNP density in winter for the S. yanoikuyae. (d) Genes with higher SNP density in summer for the M. universalis.

Functional annotation of the high-F_ST_ sets showed distinct seasonal targets in the two genera. In the genus *M. universalis*, high-F_ST_ genes were enriched in functions of cell motility (i.e. type IV pilus proteins, fimbrial protein FimT, flagellar and chemotaxis components), cross-membrane transport and cobalamin biosynthesis and transport (CobC). Additional divergent genes included stress-response and regulatory proteins (cyclic di-GMP phosphodiesterase, MarR family regulator, nitric-oxide dioxygenase, HflC and HflK) and prophage or secretion-related proteins (DNA invertase, HlyD, zona occludens toxin). KEGG pathway tallies for genes with significantly higher SNP density in summer in this genus were enriched in transporters, two-component systems, transcription factors, chromosome-associated proteins, sulfur metabolism, DNA repair and recombination, and secretion systems (Fig. 4c).

In the *S. yanoikuyae*, high-F_ST_ genes were concentrated in exopolysaccharide and surface-structure biosynthesis (polyisoprenyl-phosphate glycosyltransferase, VpsM, ExoW, ExoO, ExoZ) and flagellar assembly (FliQ, FliN, FlgC). KEGG enrichment for genes with significantly higher SNP density in winter in this genus highlighted transporters along with pathways linked to amino sugar and nucleotide sugar metabolism, two-component systems, secretion systems, peptidoglycan biosynthesis and degradation, transcription factors, and DNA repair and recombination (Fig. 4d). The F_ST_ analysis also indicated significant divergence for some genes in *Thalassarchaeum*. For instance, an ATP-binding protein exhibited higher nucleotide diversity in summer (0.0036) but lower diversity in winter (0.0008), resulting in a significantly high F_ST_ value (Fig. S7). Conversely, an ATP-binding cassette domain- containing protein showed higher nucleotide diversity in winter (0.00120934) (Fig. S6). Furthermore, in Arctic96AD-7, the “cell division protein FtsA” was identified as one such significantly divergent gene (Fig. S8).

### The M. universalis and S. yanoikuyae pangenomes reveal a pool of season-specific accessory genes

To further investigate the adaptive mechanisms at the strain level, we reconstructed the pangenomes for the two seasonally dominant species, *M. universalis* and *S. yanoikuyae*, and evaluated the differential coverage of genes between seasons. For *M. universalis*, nine high-quality MAGs (average completeness: 87.93%) were used to construct a pangenome containing 4,894 non-redundant genes, of which 1,096 formed the core genome. By mapping metagenomic reads to this pangenome, we identified 15 genes that were exclusively detected in summer samples, the majority of which were present in at least half of the analyzed MAGs. These summer-specific accessory genes primarily involved specialized transport and defense systems, including heme exporter protein D, fluoride exporters, and sulfonate transport system ATP-binding proteins, as well as prokaryotic defense components such as CRISPR-associated protein Cas2 and toxin-antitoxin (TA) systems (e.g., *parD1/3/4* and *vapB*) (Fig. 5). Similarly, for the *S. yanoikuyae* pangenome, 12 MAGs (ANI > 95%; average completeness: 89.88%) were analyzed to examine gene content variations. In contrast to *M. universalis*, the *S. yanoikuyae* population possessed a distinct set of winter-specific accessory genes. These included ribosomal proteins (e.g., *rpsU* and the large subunit cluster *rpmAEGH*), restriction-modification (R-M) systems (e.g., *mcrA*), and an extensive repertoire of TA systems (e.g., *relE*, *ccdA*, *parE1/3/4*, *doc*, and *higAB*) (Supplementary Table 5). Additionally, winter-specific genes related to transcriptional regulation, pilus assembly (*flp*), and methyl-accepting chemotaxis proteins were identified. Collectively, these results suggest that both species maintain a flexible accessory gene pool that facilitates seasonal dominance through specialized strategies in nutrient acquisition, environmental sensing, and genomic defense.

**Fig 5.**
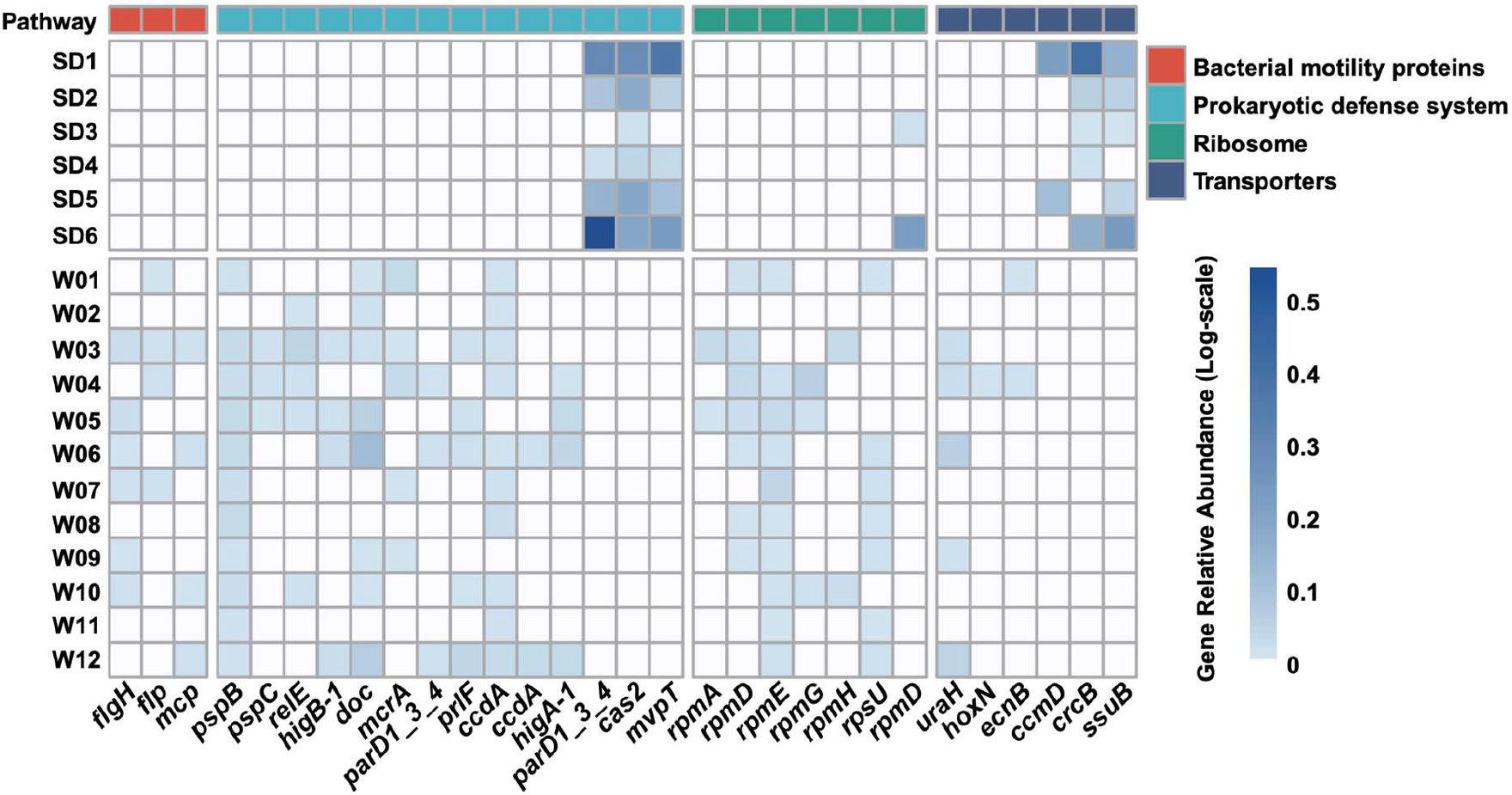
Comparative heatmap of season-specific functional genes in *S. yanoikuyae* and *M. universalis*. The heatmap illustrates genes exhibiting significant seasonal variation between summer (SD1–SD6) and winter (W01–W12) samples. Rows represent individual samples, while columns represent specific genes grouped by functional pathways (indicated by the top color bar). Color intensity reflects the log-transformed relative abundance of each gene. Gene abbreviations are detailed in Supplementary Table 6.

## Discussion

### Winter-summer differences extend to population-genomic structure

In this study, we report the *in situ* community structures varying between the seasons in the deep-sea water of SCS, which aligns with our previous study^20^. Similar winter-summer or temporal microbial changes have been reported in other ocean time-series studies and are often linked to stratification, vertical mixing, and particle flux^27–31^. By focusing on persistent microbial populations present in both sampling periods, we detected seasonal changes in within-population genomic structure. This divergence is not limited to community composition but is accompanied by systematic shifts in intra-population genetic diversity, recombination patterns, and gene content. Previous studies primarily focused on taxonomic composition^32–34^, whereas our results showed that deep-sea microbial differences between winter and summer samples can extend to the population-genomic level.

### Abundance–activity decoupling indicates distinct functional states

Differences in community composition and transcriptional activity indicate that deep-sea microorganisms respond to winter and summer environmental regimes in a taxon-specific manner. *M.universalis* was enriched in summer samples and showed transcriptional activity. In contrast, *S. yanoikuyae* was enriched in winter samples but showed no detectable transcriptional signal. Similar decoupling between abundance and activity has been reported in marine and other microbial ecosystems^35–37^. This pattern often reflects the coexistence of active, low-activity, and dormant microbial populations within the same environment^38–40^. In deep-sea environments, this decoupling may reflect temporal variation in organic matter inputs, where some taxa become active only when available substrates match their metabolic potential. The asynchronous patterns observed for *M. universalis* and *S. yanoikuyae* therefore suggest that metagenomic abundance alone may not fully capture the functional state of dominant populations.

### Enrichment without evidence for simple clonal sweeps

Enriched populations tended to exhibit higher SNP counts or SNP densities during the period in which they were more abundant, a pattern inconsistent with a simple clonal sweep. Rapid clonal sweeps usually reduce within-population diversity, whereas the coexistence of high abundance and high diversity points to the expansion of genetically heterogeneous populations composed of multiple closely related strains. In this study, both *M. universalis* and *S. yanoikuyae* showed elevated SNP levels under the winter or summer conditions in which they were enriched. This pattern suggests that their enrichment did not arise from the replacement by a single low-diversity genotype. This interpretation is consistent with previous observations that microbial ecological success can involve diverse strain assemblages rather than genome-wide selective sweeps^11,41,42^. Such intra-population variation may provide a reservoir of allelic diversity that helps deep-sea microorganisms respond to fluctuating environmental conditions.

### Summer-enriched M. universalis shows resource-exploration features

Abundant marine *Prochlorococcus* populations can contain multiple ecologically differentiated subpopulations^43^. High nucleotide diversity alongside gene-specific sweeps has also been reported in subseafloor aquifer populations^44^. Similarly, the summer-enriched *M. universalis* population was characterized by increased abundance, elevated nucleotide diversity, and stronger divergence in genes related to motility, chemotaxis, and nutrient uptake. This combination is difficult to reconcile with a recent genome-wide sweep, which would be expected to reduce within-population diversity^28^. Instead, the summer-enriched *M. universalis* is better interpreted as a heterogeneous assemblage with divergence concentrated in niche-related functions. In marine environments, motility and chemotaxis are central to bacterial responses to nutrient patchiness, particle-associated habitats, and transient organic matter hotspots^45–47^. More broadly, bacterial diversification often concentrates in genes involved in environmental sensing^48–50^, surface interaction^51,52^, and substrate transport^53,54^. Summer-associated accessory genes further suggest that enrichment of *M. universalis* involved not only allelic shifts but also flexible gene content^55^.

F_ST_ analysis further showed that winter-summer differentiation in *M. universalis* was concentrated in specific functional modules rather than distributed uniformly across the genome. High-divergence genes were associated with cell-surface systems, type IV pili, motility, chemotaxis, transport, regulation, and cobalamin-related pathways. Divergent type IV pilus and flagellar genes may facilitate particle colonization^56,57^, surface attachment^58^, and resource acquisition^59^. Higher summer surface productivity in the SCS may increase organic matter and nutrient inputs to mesopelagic depths, providing a plausible environmental context for the observed divergence in motility, chemotaxis, and transport-related genes. Therefore, the summer-associated genomic profile of *M. universalis* is consistent with niche exploration and resource acquisition during the sampled summer period, rather than simple expansion of a single lineage.

### Winter-enriched *S. yanoikuyae* shows a defense-oriented persistence profile

During winter in the northern South China Sea, monsoon-driven mixing can weaken stratification and enhance vertical exchange^60,61^, potentially altering particle fluxes and trace-metal inputs to deeper waters^62^. Within this environmental context, high-Fst genes in the winter-enriched *S. yanoikuyae* population revealed a distinct set of genomic features. Many divergent genes were involved in exopolysaccharide (EPS) and surface-structure biosynthesis, including polyisoprenyl phosphate glycosyltransferase, VpsM, ExoW, ExoO, ExoZ. These EPS and surface-structure genes may support attachment, aggregation, and persistence under resource-limited or physically variable conditions^63^. In the winter-enriched *S. yanoikuyae* population, metal resistance and detoxification genes, including CopB, CopC, multicopper oxidases, Cu^2+^-ATPase, and CnrH showed elevated diversity and divergence. These genes may complement EPS-mediated metal sequestration and potentially increase tolerance to metal-related stress. Metal-related stress, particularly from copper and iron, can impose ecological pressure in oligotrophic marine ecosystems^64^. It may become more relevant during winter mixing, when vertical exchange can alter particle fluxes and trace-metal inputs to deeper waters^65,66^.

Winter-associated divergence was also observed in genes related to DNA replication, protein stability, and oxidative stress, including alcohol dehydrogenase, aldehyde dehydrogenase, and Hsp15. These genes may contribute to cellular maintenance by mitigating oxidative damage and maintaining protein integrity under stressful conditions. Such functions may help *S. yanoikuyae* cope with increased exposure to trace metals and complex organic matter during intensified vertical mixing^67,68^. These stressors can also reduce microbial metabolic activity^69^. Similar stress-response patterns have been reported in cold-adapted bacteria under low-temperature conditions^70^. Together, these features are consistent with a persistence-oriented profile in the winter-enriched *S. yanoikuyae* population.

Viral predation is a major evolutionary force shaping microbial genome architecture and defense systems in marine environments. Similar microbe-virus arms races have also been reported in deep-sea environments^71^. In the winter-enriched *S. yanoikuyae* population, winter-associated accessory genes included restriction-modification systems and toxin-antitoxin modules, including *relE*, *ccdA*, *parE*, and *higAB*. These defense-related genes are consistent with protection and persistence-associated functions under oligotrophic conditions^72,73^. Toxin-antitoxin systems have also been linked to persister-cell formation and survival during prolonged stress or starvation^74,75^. This interpretation is consistent with the lack of detectable transcripts from *S. yanoikuyae* despite its winter enrichment. Winter-associated ribosomal proteins and transcriptional regulators suggest that this population may retain functional capacity under low-activity conditions^76^. Chemotaxis-related genes may further support response to episodic resource availability in the deep sea^77^. Together, these features support a defense-oriented persistence profile in the winter-enriched *S. yanoikuyae* population.

## Conclusion

This study reveals the deep pelagic ocean as a dynamic ecosystem shaped by predictable seasonal evolutionary processes. Through metagenomics, metatranscriptomics and population genomics, we uncover a pronounced succession between two key microbial lineages employing contrasting strategies: the opportunistic, growth-focused *M. universalis* dominant in summer, and the defense-oriented, survival-focused *S. yanoikuyae* dominant in winter. These divergent genomic adaptations, influenced by selection, recombination, and flexible pangenomes, drive seasonal shifts in deep-sea biogeochemical cycling, impacting organic matter remineralization and trace element dynamics.

Our results highlight the central role of microbial evolutionary dynamics in shaping ecosystem function and suggest that viral pressures further complicate these seasonal patterns. Integrating metaproteomics will be essential to confirm the functional expression of these adaptations. Understanding these lineage-specific strategies is critical for predicting deep-sea ecosystem resilience to climate change–induced alterations in seasonal environmental pulses.

## Methods

### Deep-sea water column sampling and nucleic acid extraction

The sampling was performed during two expeditions conducted from November 23-25, 2019 (winter) and June 9-11, 2020 (summer). Samples were collected from depths ranging from 1,000 to 1,100 meters in the South China Sea using the MISNAC system on the ‘Phoenix’ lander^22^ (Fig. S1). During the winter expedition, two replicates were collected each day over a 24-hour period, yielding 12 samples from six MISNAC subunits, with each sampling session lasting approximately four hours. More details of the sampling have been described in our previous publications^20,21^.

### Metagenomic sequencing and quality control

For DNA library preparation, the 18 MISNAC samples were incubated with 0.1 ng/μl RNaseA (TaKaRa, Dalian, China) at 37 ℃ for 10 min to remove RNA fragments. Approximately 100 ng of DNA was fragmented to an average size of 350 bp by ultrasouication using a Covaris M220 Focused ultrasonicator (Covaris, Massachusetts, USA), and high-throughput metagenomic libraries were subsequently constructed using the VAHTS Universal DNA library prep kit for Illumina V3 (Catalog No.: N411-01/02/03, Vazyme, Nanjing, China) according to the manufacturer’s protocol. The libraries were sequenced on an Illumina NovaSeq6000 platform (Illumina, San Diego, USA) using paired-end 2 × 150 bp sequencing.

Raw Illumina reads were trimmed to remove adapters and filtered using fastp^78^(v0.21.0) with the following parameters: (-f 1 -t 0 -F 1 -T 0 -w 24 -c -q 20 -u 20 -g -W 5 -3 -l 50). Reads with a quality score below 20 for more than 20% of their length, unpaired reads and reads shorter than 50 bp were removed. Duplicated paired Illumina reads were removed by FastUniq^79^ with default parameters. Illumina metagenomic reads were mapped to a contaminant database (including mouse, human and common laboratory contaminant bacterial genomes downloaded from NCBI) using Bowtie2^80^ with default parameters to filter out contaminant reads.

### Metagenomic assembly and genome binning

Each of the 18 Illumina datasets was assembled individually using metaSPAdes (v3.15.2)^81^ with the parameters ‘-min-contig-len 1300 -k 21,33,55,77,99,127’. For the assembled contigs, MAGs were binned separately from the winter and summer assemblies using MaxBin^82^, MetaBAT^83^, and CONCOCT^84^ as integrated in MetaWRAP (v1.2.1)^85^. The bin refinement module was run using default parameters. MAGs with >50% completeness and < 10% contamination (metawrap bin_refinement ‘-c 50 -× 10’) were identified and using CheckM^86^. Dereplication of these MAGs was performed using the dRep (v3.5.0)^87^ with an ANI threshold of 95% at species level (- pa 0.95 -sa 0.99 -nc 0.30 -cm larger -p 40 -comp 50 -con 10). As the result, 426 MAGs were gained for further analysis.

### SNP identification and F_ST_ calculation

All the MAGs were combined to create a reference genome database against which paired-end reads (150 bp by 2) from each sample were mapped with Bowtie2 in sensitive mode. Genome-wide inStrain^88^ profiles were conducted with default settings. Relative abundances of MAGs in function of RPKM (Reads Per Kilobase per Million mapped reads) were determined using the genome-wide average read mapping coverage, which was conducted by CoverM, using the rpkm mode. Only MAGs with an average coverage of > 5× and breadth (fraction of the MAG covered by at least one read of a metagenome) of >0.5 (at least 50% of the MAG could be covered by the metagenomes for a season) were included for relative abundance calculation and further analyses. inStrain profile was used to call SNPs and calculate nucleotide diversity. pN/pS ratios of genes were calculated for each MAG in a sample by inStrain profile and were averaged for the calculation of the genome-wide pN/pS ratio. Linkage disequilibrium (r^2^) was calculated for all pairs of SNPs with at least 20 pairs of connecting reads and normalized by rarefying to 20 read pairs using inStrain. The F_ST_ (fixation index) (measure of differential allele frequencies between populations) value for each gene of all populations were calculated between summer and winter metagenomic datasets using a method previously described by Crits-Christoph et al. ^89^.

### Pangenome analysis

For the pangenome construction, we used Roary v3.13.0^90^ with flag -s on quality-filtered (completeness of >70% and contamination of <10%) MAGs binned from metagenomic samples and clustered with representative *M. universalis* at >95% ANI. All genes in the pangenome were extracted to create a reference database to which all reads across all metagenomic samples from winter and summer were mapped using Bowtie2 in a sensitive mode and filtering for a maximum of 15 mismatches (>90% sequence identity) per read. Relative abundance of genes was determined by normalizing the coverages of genes by the average coverage of representative *M. universalis* MAGs in each sample. The genes with relative abundance of >3 were discarded as they are likely also present in populations other than *M. universalis*^28^. Variation in relative gene coverage in all samples from summer versus winter was assessed via ANOVA, and the resulting *p* values were adjusted for multiple-hypothesis tests using the Benjamini-Hochberg procedure. Genes with an adjusted *p* value of <0.01 were classified as season specific if present with a coverage of <10% of the average coverage of *M. universalis* MAGs across all time intervals in either summer or winter.

### Statistical analyses

All statistical tests were performed using R (v4.3.2)^91^ and visualized using ggplot2^92^.

## Acknowledgments

This study was supported by National Natural Science Foundation of China (42376149) and Shenzhen Key Laboratory of Advanced Technology for Marine Ecology (ZDSYS20230626091459009).

## Founding statement

This study was supported by the National Natural Science Foundation of China (Grant No. 42376149) and the Shenzhen Key Laboratory of Advanced Technology for Marine Ecology (Grant No. ZDSYS20230626091459009).

## Ethics statement

No specific ethical approval was required for this study as it did not involve humans, endangered or protected species, or laboratory animals.

## Data availability

All data are available in the main text or the supplementary materials. The raw metatranscriptome sequencing data, assembled contigs and MAG sequences generated in this study have been deposited in the NCBI database under the accession code PRJNA1067282.

---

**Fig. S1.**
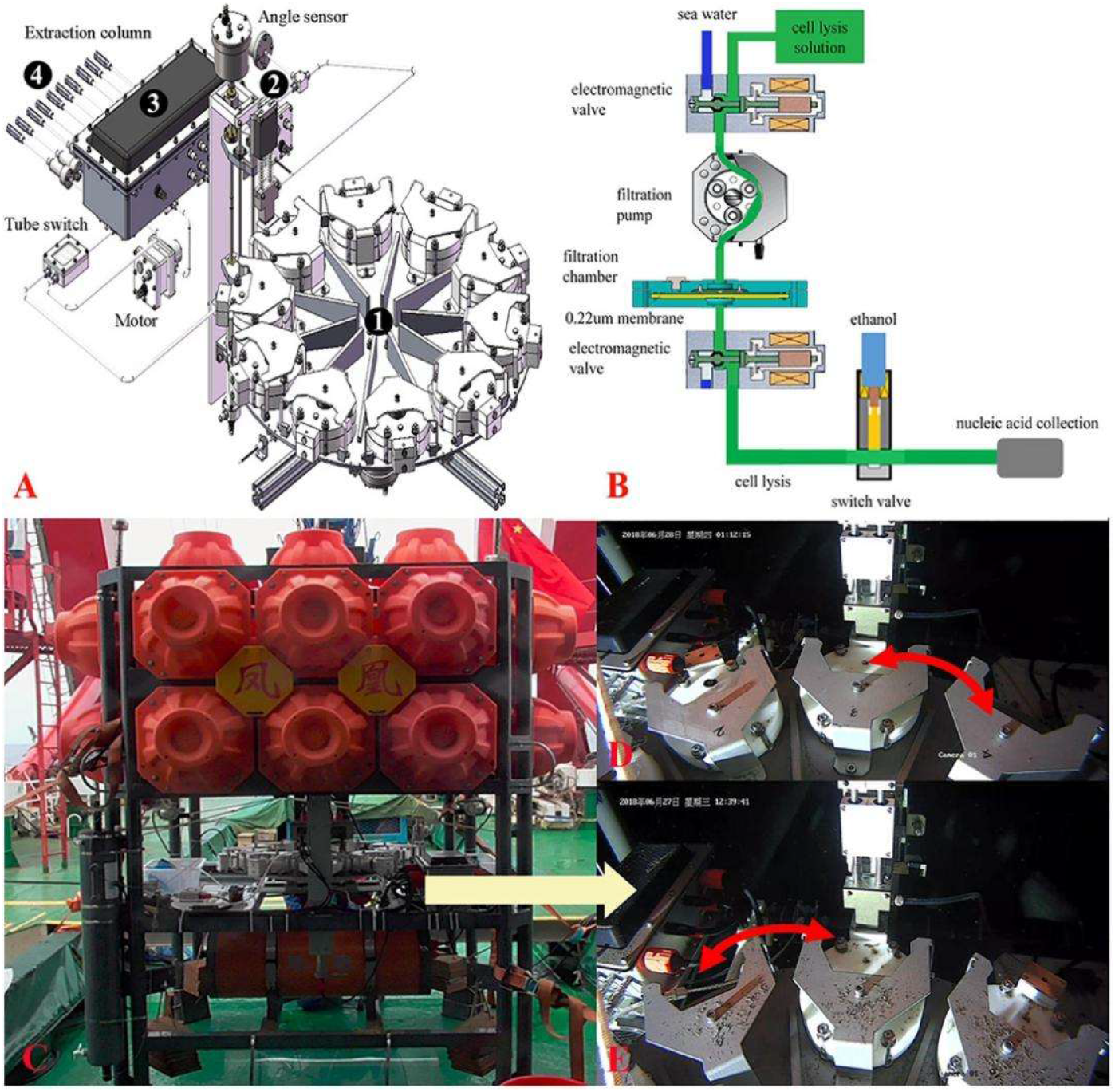
Schematic structure of multiple in situ nucleic acid collections (MISNAC) from deep-sea zone. The four parts of MISNAC: a chamber supporting and transition subsystem (1), a pipe docking subsystem (2), a flow control subsystem (3) and a nucleic acid collection subsystem (4) are labeled on the diagram (A). The workflow of nucleic acid extraction was demonstrated in (B). The MISNAC apparatus was equipped on a lander named ‘Phoenix’ (C). The switch of filtration chambers of the MISNAC apparatus was demonstrated by the photographs taken by a camera on the lander (D,E).

**Fig. S2.**
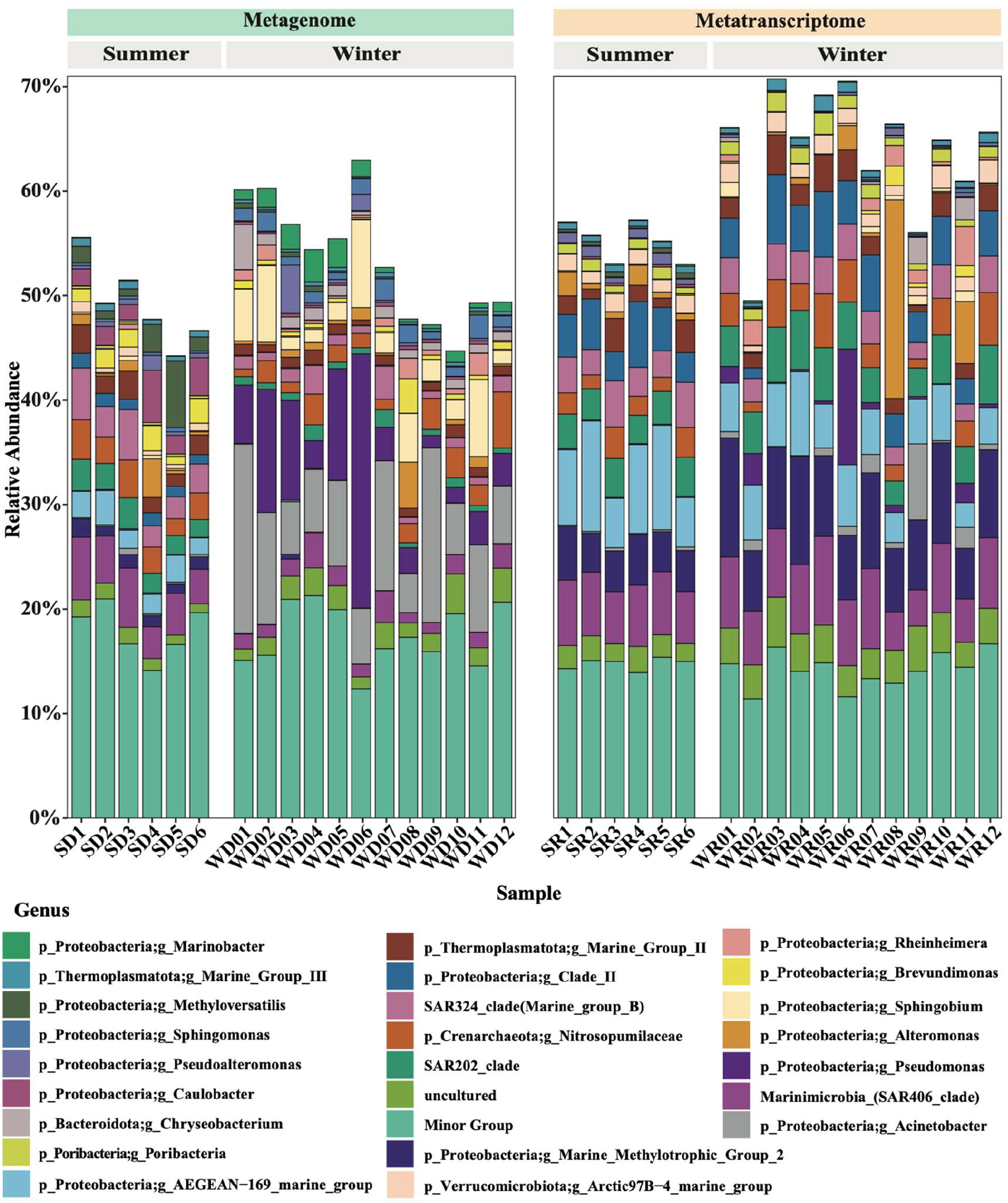
Seasonal variation of the community structures at the sampling sites. Taxonomic composition of prokaryotic community of metagenomes and metatranscripomes at genus level based on the classification of 16S rRNA gene sequences.

**Fig. S3.**
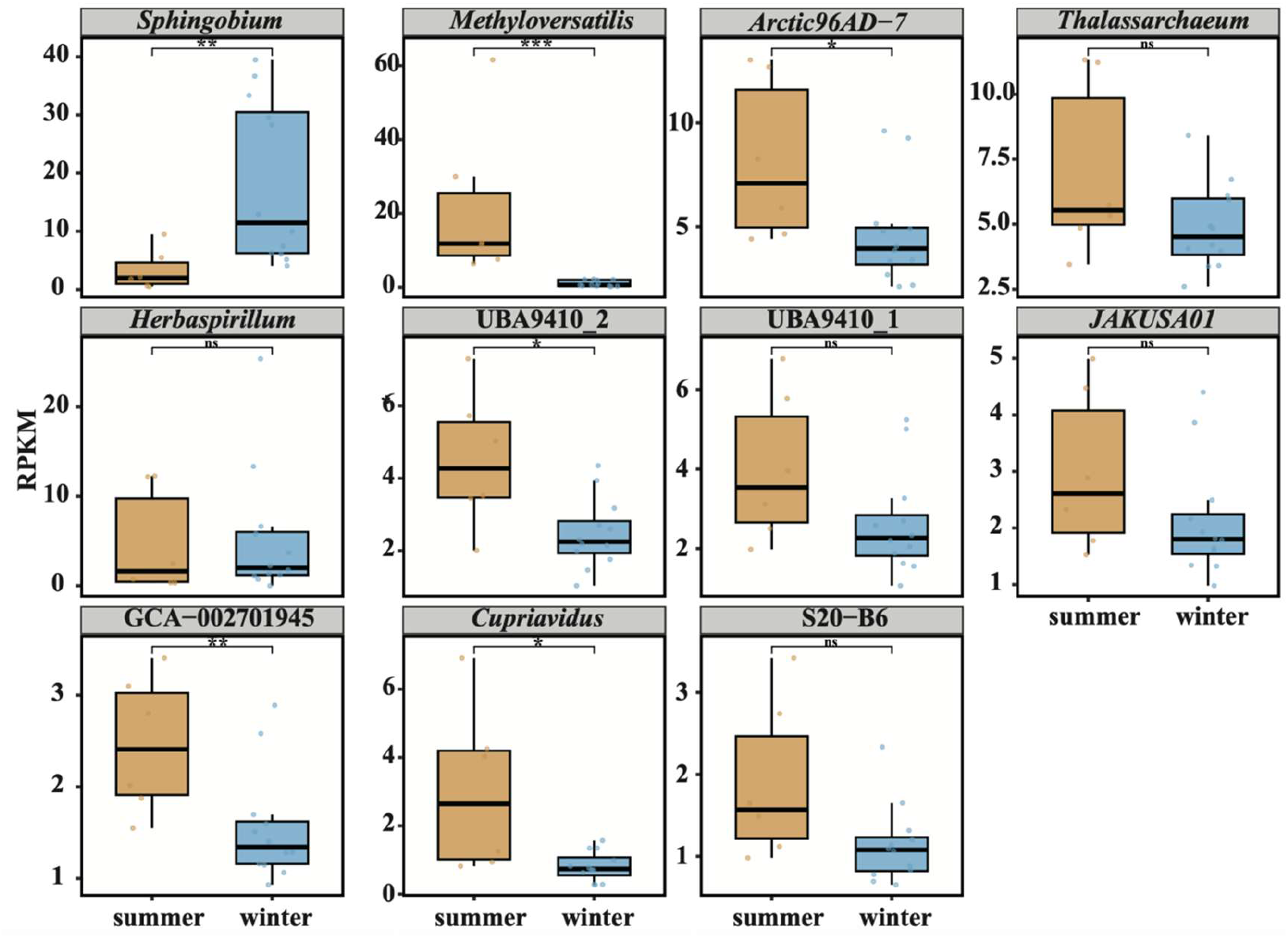
Seasonal differences in abundance (RPKM) of MAGs representing 11 species. Boxplots compare the distribution of RPKM values between summer (orange) and winter (blue) for each genus. Asterisks indicate statistically significant differences (*p* < 0.05, *p* < 0.01, *p* < 0.001; Wilcoxon rank-sum test). *Methyloversatilis universalis* and GCA-002701945 were significantly more abundant in summer, whereas *Sphingobium yanoikuyae* was enriched in winter. Several other genera showed season-specific abundance trends but without statistical significance.

**Fig. S4.**
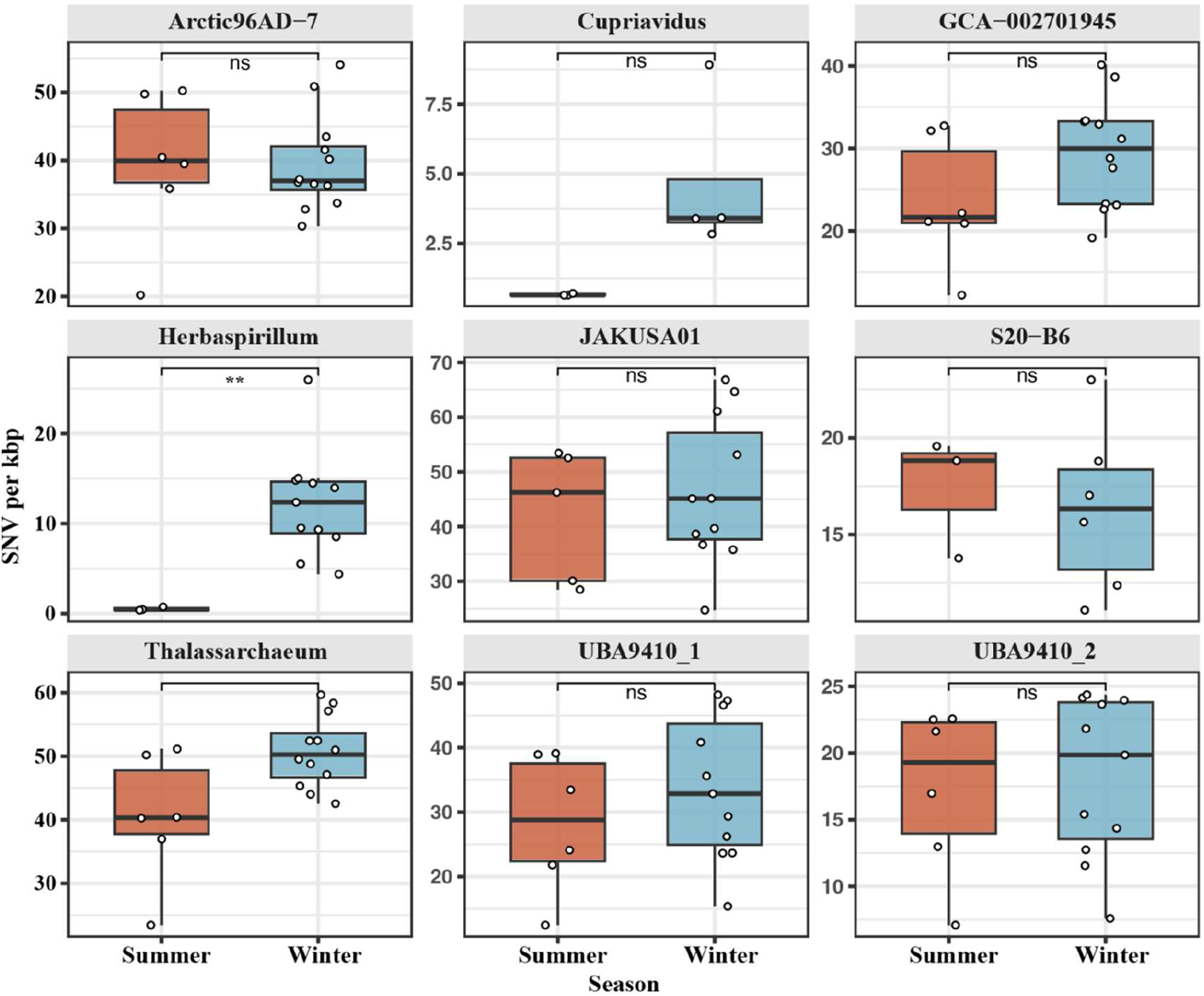
Comparison of single nucleotide variation density between summer and winter samples across another 9 selected genera. Boxplot comparison of SNV densities between summer and winter for the 9 species. Asterisks (**) indicate significant differences (Wilcoxon rank-sum test, *p* < 0.01).

**Fig. S5.**
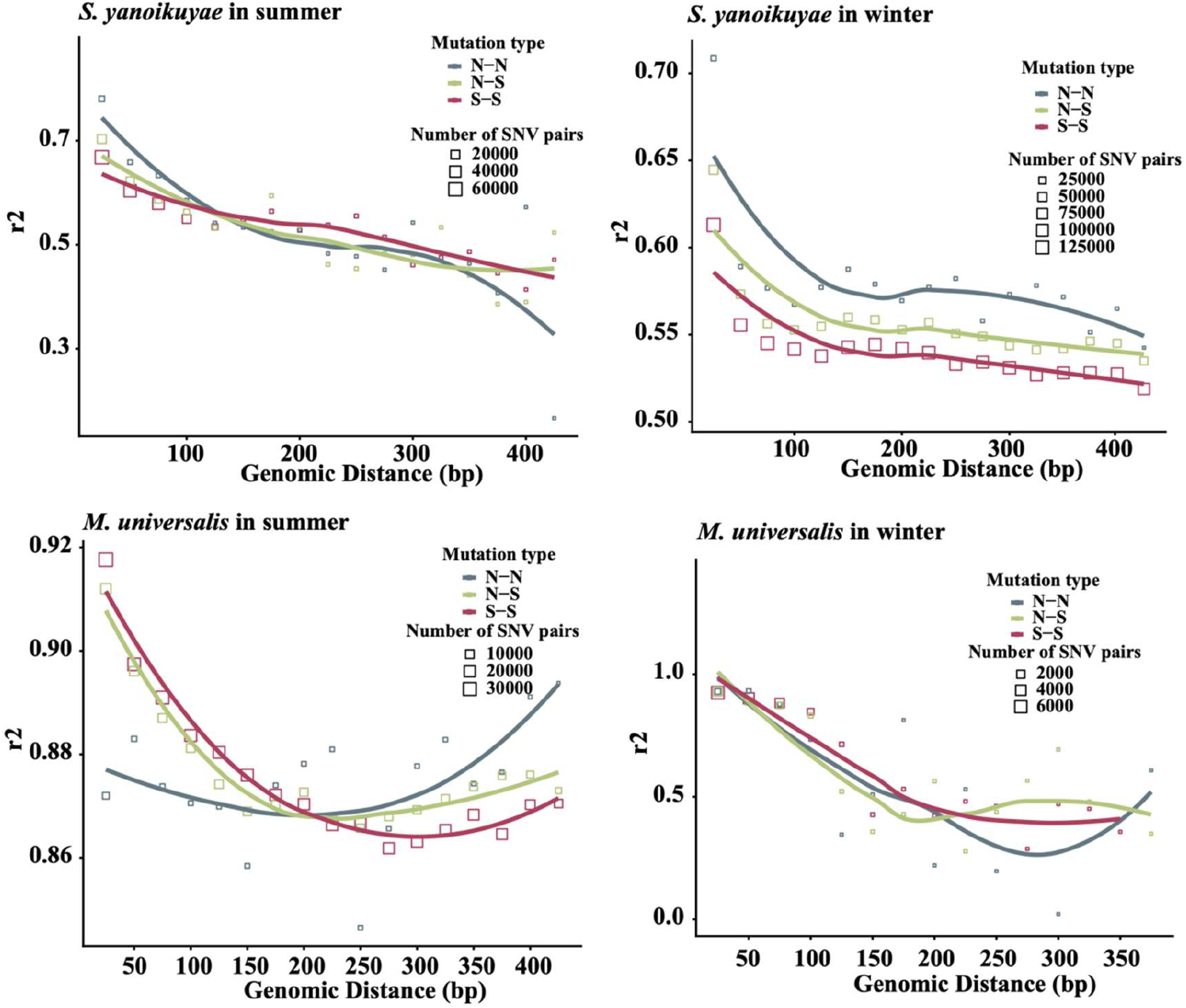
Linkage disequilibrium (LD) decay in the dominant seasonal populations *Sphingobium yanoikuyae* and *Methyloversatilis universalis*. The plots show the decay of LD, measured by the squared correlation coefficient (r²), as a function of genomic distance (bp) between pairs of single nucleotide variants (SNVs). The analysis is shown for the *Sphingobium* population in (a) winter and (b) summer, and for the *Methyloversatilis* population in (c) summer and (d) winter. Lines are fitted to show the decay trend for different mutation types: nonsynonymous-to- nonsynonymous (N-N), synonymous-to-synonymous (S-S), and mixed nonsynonymous-synonymous (N-S) pairs. The overall decay of r² with distance in all panels is characteristic of active homologous recombination. The consistently stronger linkage (higher r²) for N-N pairs compared to S-S pairs is a genomic signature indicative of recent selective sweeps targeting functional variants.

**Fig. S6.**
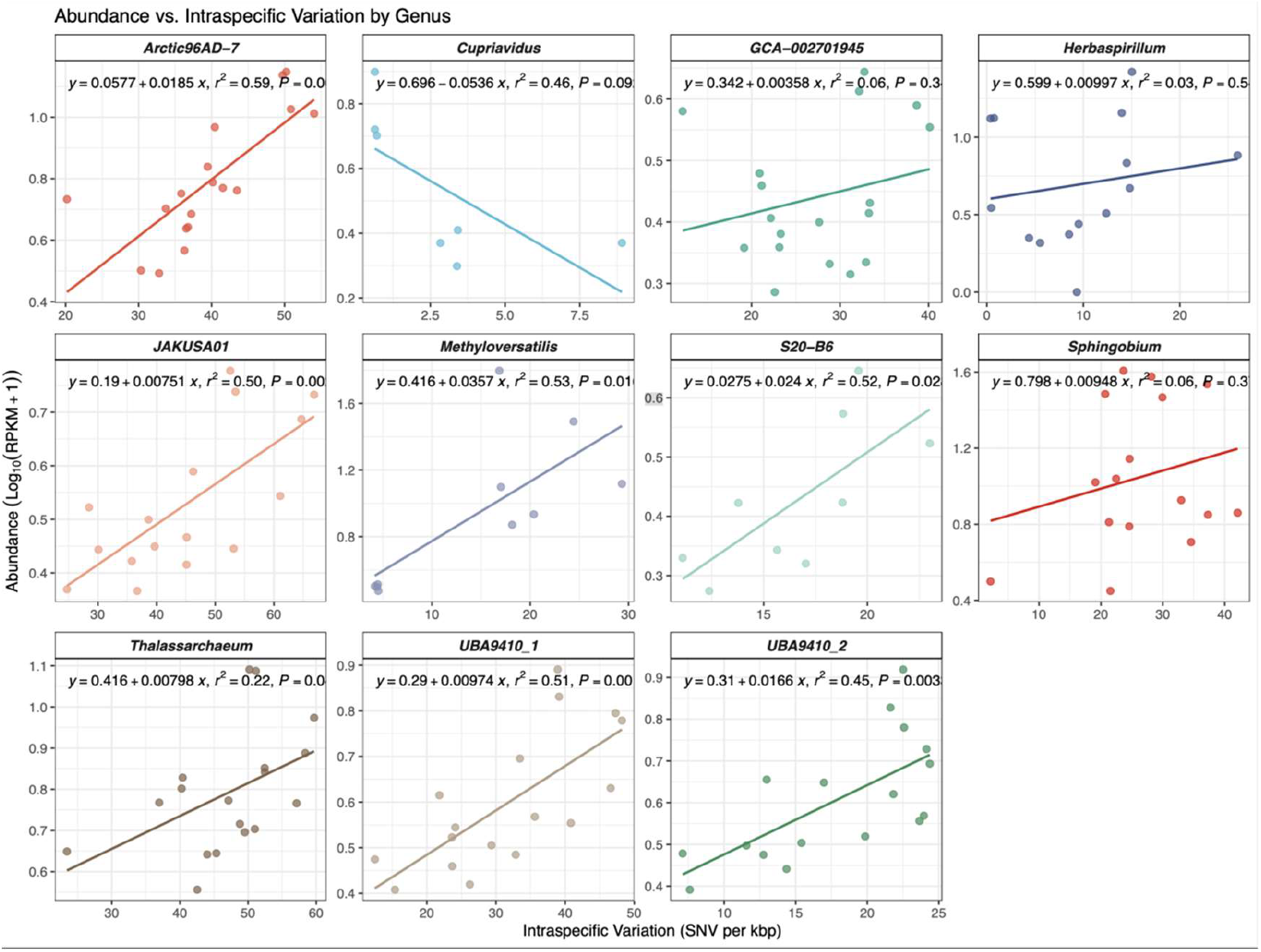
Correlation between population abundance and intraspecific genomic diversity across the studied deep-sea microbial species. The scatter plots illustrate the relationship between the relative abundance (log-transformed RPKM) and the density of single nucleotide variants (SNVs per kbp) for the major genera identified in the deep-sea metagenomic samples. Each data point represents an individual sample.

**Fig. S7.**
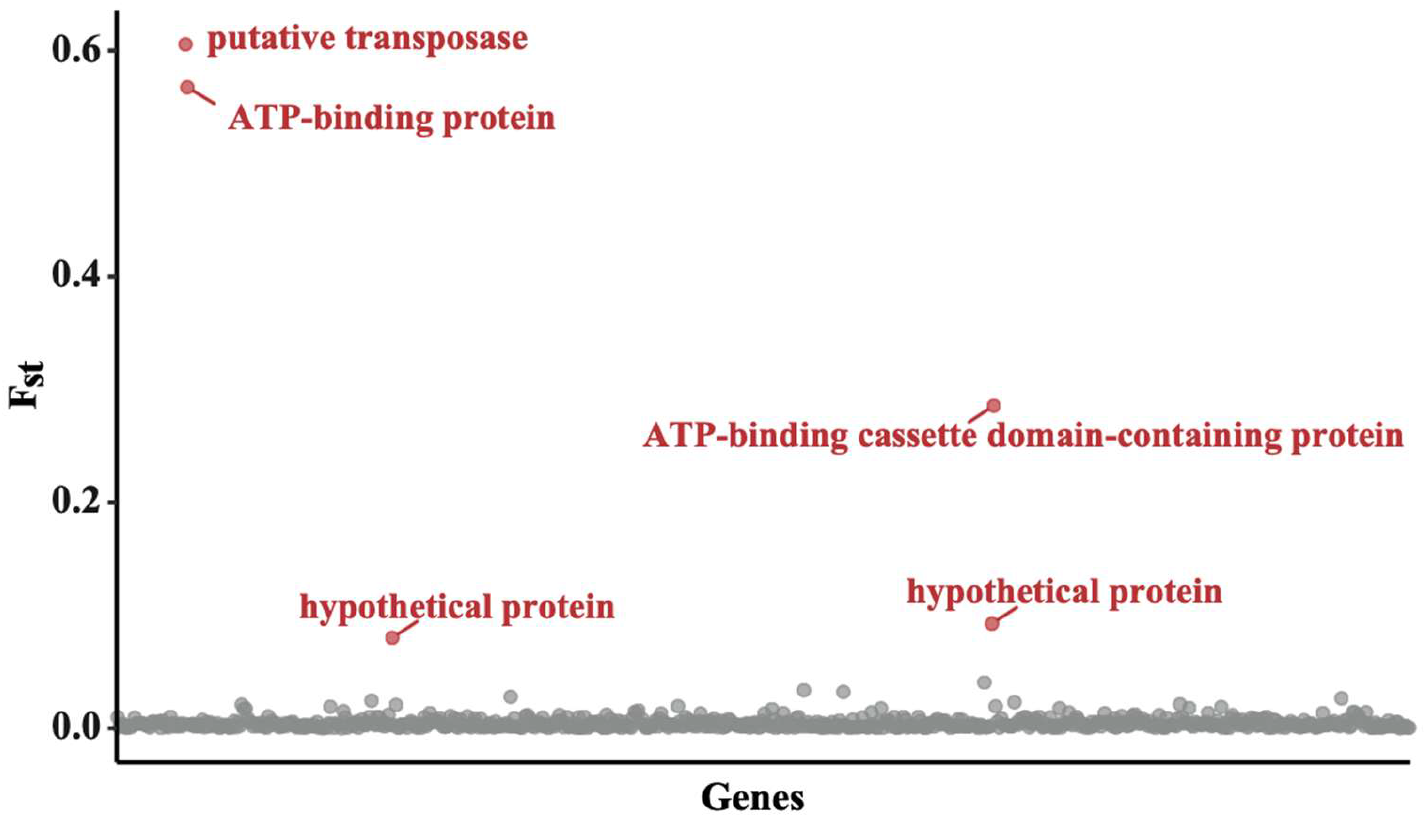
Differentiated genes (marked in red) with elevated F_ST_ values with statistical significance (adjusted *p* value of < 0.05 [by a Z test]) from *Thalassarchaeum*.

**Fig. S8.**
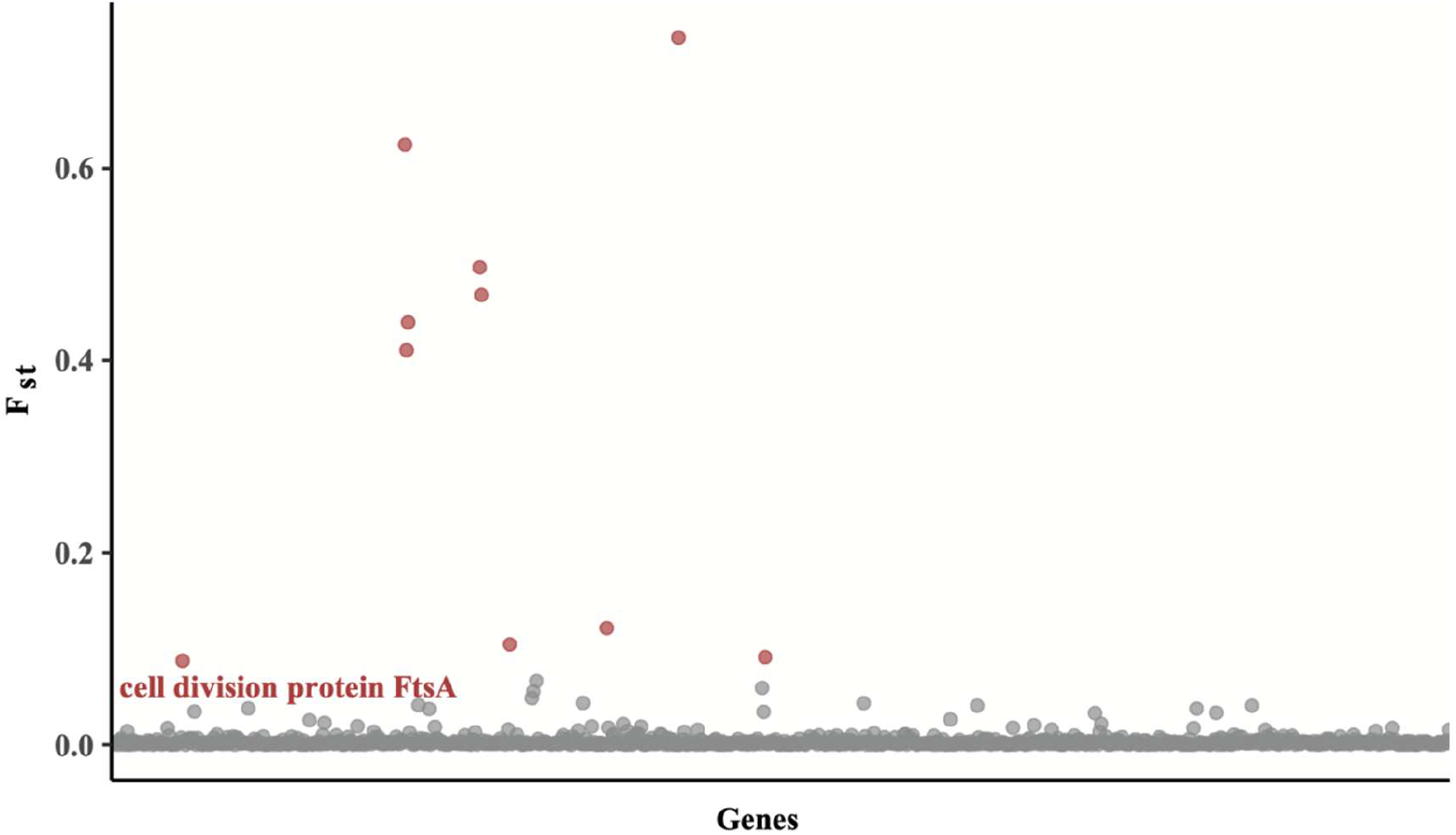
Differentiated genes (marked in red) with elevated F_ST_ values with statistical significance (adjusted *p* value of < 0.05 [by a Z test]) from Arctic96AD-7.

